# Selective sweeps in populations of the broad host range plant pathogenic fungus *Sclerotinia sclerotiorum*

**DOI:** 10.1101/352930

**Authors:** Mark C Derbyshire, Matthew Denton-Giles, James K Hane, Steven Chang, Mahsa Mousavi-Derazmahalleh, Sylvain Raffaele, Lone Buchwaldt, Lars G Kamphuis

## Abstract

The pathogenic fungus *Sclerotinia sclerotiorum* infects over 600 species of plant. It is present in numerous environments throughout the world and causes significant damage to many agricultural crops. Fragmentation and lack of gene flow between populations may lead to population sub-structure. Within discrete recombining populations, positive selection may lead to a ‘selective sweep’. This is characterised by an increase in frequency of a favourable allele leading to reduction in genotypic diversity in a localised genomic region due to the phenomenon of genetic hitchhiking.

We aimed to assess whether isolates of *S. sclerotiorum* from around the world formed genotypic clusters associated with geographical origin and to determine whether signatures of population-specific positive selection could be detected. To do this, we sequenced the genomes of 25 isolates of *S. sclerotiorum* collected from four different continents - Australia, Africa (north and south), Europe and North America (Canada and the northen United States) and conducted SNP based analyses of population structure and selective sweeps.

Among the 25 isolates, there was evidence for four population clusters. One of these consisted of 11 isolates from Canada, the USA and France (population 1), another consisted of five isolates from Australia and one from Morocco (population 2). A further cluster was made up of Australian isolates, and the single South African isolate appeared to be from a separate population. We found that there was evidence of distinct selective sweeps between population 1 and population 2. Many of these sweeps overlapped genes involved in transcriptional regulation, such as transcription factors. It is possible that distinct populations of *S. sclerotiorum* from differing global environments have undergone selective sweeps at different genomic loci. This study lays the foundation for further work into investigation of the differing selective pressures that *S. sclerotiorum* populations are subjected to on a global scale.

## Introduction

Spread of a favourable allele through a population due to positive selective pressure is known as a selective sweep. The increase in frequency of a single favourable allele toward fixation is known as a hard selective sweep. On the other hand, the increase in frequency of multiple adaptive alleles conferring equivalent fitness benefits is known as a soft selective sweep (1,2). When a favourable allele increases in frequency due to positive selective pressure, linked loci will also increase in frequency. Linkage of non-selected loci decreases with distance from the favourable allele due to the effects of recombination (3). Thus, immediately following a hard selective sweep, the genomic region surrounding the beneficial locus will be largely monomorphic throughout the population.

Several tests for selective sweeps have been developed based on this understanding (4–6). Such tests have been applied to numerous organisms to study the footprints of positive selection in genomes (7). In most studies, evidence of selective sweeps localised to particular populations has been observed. For example, 346 genomic regions in *Drosophila melanogaster* were found to exhibit a significantly high degree of fixation between populations from Europe and Africa. Genes within these regions were enriched for various functional activities including transcriptional regulation (8). In humans, numerous population specific signals of positive selection have been identified and linked with genes involved in processes such as skin pigmentation, immunity, heat shock and olfactory perception (9–12).

Plant pathogenic fungi may be useful organisms for the study of positive selection. This is because they likely undergo rapid adaptation to anthropogenic selective pressures such as dissemination to naive environments and introduction of new crop varieties or antifungal chemistries (13–20). A selective sweep scan of a fungal plant pathogen was recently published. This study identified differential selective sweeps between two species of *Microbotryum*. Several of the genes associated with selective sweeps in these species exhibited domains and expression patterns indicative of roles in interaction with the host (21). A further preprint article is also available detailing selective sweeps in the plant pathogenic fungus *Rhynchosporium commune*. In this study it was found that numerous, non-overlapping sweeps appeared in different populations. The kinds of genes associated with selective sweeps encoded proteins with functions related to biotic and abiotic stress response (22).

To gain further insight into positive selective pressure in a plant pathogen, we examined the genome of the fungus *Sclerotinia sclerotiorum* for evidence of selective sweeps. The genome sequence of *S. sclerotiorum* was recently sequenced to completion using PacBio technology and has been annotated using extensive RNA sequencing data and manual curation (23). This species is a host generalist that infects more than 600 species of plant (24). It has been globally disseminated and is present in numerous environments where it causes significant yield losses in economically important crops (25). Several studies have demonstrated genotypic differentiation of *S. sclerotiorum* populations from different geographic regions. For example, Kamvar et al. showed that populations of *S. sclerotiorum* from Mexico shared no multilocus haplotypes with populations from the United States, Australia and France (26). Similarly, Lehner et al. showed that 29 % of variance within a sample of isolates from southeastern Brazil and the United States was explained by differences between populations from the two regions (27). And, echoing these findings, Clarkson et al. found that populations of *S. sclerotiorum* from the UK, Norway and Australia shared no multilocus haplotypes (28).

To identify which genomic regions may be under selective pressure in different populations of *S. sclerotiorum*, we sequenced the genomes of a collection of 25 isolates from Canada, France, the USA, South Africa, Morocco and Australia. These isolates appeared to be sampled from populations that were broadly associated with climate of origin. Within two of these populations, distinct selective sweeps were identified in different genomic regions. Many of these sweeps contained genes involved in transcriptional regulation, such as transcription factors, methyltransferases and protein kinases. We speculate that adaptation to contrasting environments has been partly mediated by changes in gene regulation in *S. sclerotiorum*.

## Methods

### Fungal isolates

In this study a total of 26 isolates were analysed, including the reference isolate, for which a continuous chromosome sequence is available (NCBI Bioproject ID: PRJNA348385) (23). These include isolates from Australia (× 12), Europe (× 5), North America (× 6) and Africa (× 2). The names of the isolates, host of origin (if known), location, year collected, accession numbers (where applicable) and names of isolate stock collections (where applicable) are summarised in Supplementary Table 1. The isolates originate from regions with a range of different climates, which include the following distinct climate types according to the Köppen climate classification scheme (29): Csa, hot summer Mediterranean; Csb, warm summer Mediterranean; Cfb, Oceanic;Dfb, humid continental.

### DNA extraction and sequencing of fungal isolates

DNA from the isolates Ss44, Ss45, Sssaf, SsChi, BloC104, BloC014, P134, P163, FrB5, SK35, 321, MB52, MB21 and AB29 was extracted and sequenced in a previous publication (30). DNA from all Australian isolates apart from CULm and CULa was extracted and sequenced using the following method: sterile sclerotia of each isolate were plated onto full strength potato dextrose agar (PDA) and grown for 4-10 days at 20 °C in the dark. Mycelium from the edge of each colony was harvested and DNA was extracted using a modified version of the alkaline lysis method described by Rollins (31). In brief, following alkaline lysis, DNA solutions were partitioned using 24:1 chloroform:isoamyl alcohol and precipitated with isopropanol and lithium chloride before being resuspended in 30 μl of milliQ water + 1 μl of RNAse A (10 mg/ml). High quality DNA was sequenced on the Illumina Hiseq platform using 2 × 250 bp paired end sequencing.

DNA from the isolates CULm and CULa was extracted and sequenced using the following method: A seven day old fungal culture grown on PDA was harvested and homogenised with pestle and mortar by adding 1 mL of half strength potato dextrose broth (PDB). The suspension was transferred to a 250 mL flask containing 100 ml of half strength PDB. The flask was shaken at 100 rpm and kept in darkness for 4 days at 20 °C. After 96 days post inoculation, the fungal culture was filtered through sterile cheese cloth and washed twice with sterile distilled H_2_O to remove any excess nutrients. The culture was freeze-dried overnight before genomic DNA extraction using the CTAB method (32). DNA concentration was quantitated using a Qubit 2.0 fluorometer (Life Technologies) and assessed by gel electrophoresis for quality control. Sequencing was performed by the Australian Genome Research Facility using the Illumina Hiseq platform with 2 × 250 bp paired end sequencing.

### *HSP60* phylogeny to assign isolates to the species *Sclerotinia sclerotiorum*

To assess whether isolates were the species *S. sclerotiorum*, a phylogenetic tree was built using previously published heat shock protein 60 (*HSP60*) gene sequences (33) in conjunction with *HSP60* gene sequences derived from *de novo* assemblies. GenBank accessions for previously published *HSP60* sequences are in Supplementary Table 2. *HSP60* sequences were extracted from *de novo* assembled contigs (*de novo* assembly of fungal isolate genomes is detailed in a later section ‘Quality filtering and *de novo* assembly of fungal isolate genomes’). Exonerate version 2.2.0 (34) was used to align the previously published *HSP60* gene sequence for *S. sclerotiorum* to each *de novo* assembly. The commands ‘--bestn 1 --mode est2genome’ were used to identify the single best alignment for each genome. The coordinates of these alignments were extracted to produce browser extensible data (BED) formatted files containing strand information. Bedtools version 2.25.0 (35) was then used with the commands ‘getfasta -s’ in conjunction with these files and the *de novo* assemblies to extract regions aligning to the published HSP60 sequence. The option ‘-s’ was used to take into account strand information of the alignments. The isolate specific and previously published *HSP60* sequences were aligned using ClustalW version 2.1 (36) with default settings. The appropriate nucleotide substitution model was chosen using JModelTest version 2.1.10 (v20160303) (37) with the settings ‘-s 11 -f -i -g 4 -AIC -BIC -AICc -DT -p -a’. The Akaike Information Criterion (AIC) test was used to determine which nucleotide substitution model to use. The model used was the ‘transitional model’ (TIM) first described in (37).

An *HSP60* based phylogeny was constructed from the alignments using PhyML version 20120412 (38) with the commands ‘-b 100 -d nt -n 1 -m 012230 -f 0.25,0.25,0.25,0.25 -c 4 -a e --no_memory_check --r_seed 12345 -o tlr -s BEST’ (with the TIM substitution model). Support for branches was assessed using 100 bootstrap tests. The species *Botrytis cinerea* and *Monilinia fructigena* were used as outgroups.

### Quality filtering and *de novo* assembly of fungal isolate genomes

Paired reads from each of the *S. sclerotiorum* isolates were assessed for adapter contamination and poor quality sequence content using FASTQC version 0.11.4 (39). Illumina data that required adapter and poor quality sequence removal were filtered using Trimmomatic version 0.36 (40); the adapter file (‘TrueSeq.fasta’) used with Trimmomatic contained sequences for the TruSeq LT and v1/v2 kits. Information on whether isolates were quality and adapter filtered is included in Supplementary Table 1. The paired Illumina reads from each isolate were *de novo* assembled using the A5-miseq pipeline version 20160825 (41) with default settings.

### Mapping of Illumina reads to the reference genome

Paired reads for each of the isolates were mapped to the reference genome using Stampy version 1.0.31 with default settings (42). Both paired reads and singletons that survived the previously described quality filtering step (see ‘Quality filtering and *de novo* assembly of fungal isolate genomes’) were mapped to the reference genome. Reads were first mapped using BWA MEM version 0.7.5a-r405 (43) and resulting binary alignment map (BAM) files were used as input to Stampy.

### Variant calling and quality filtering

Following alignment, the Genome Analysis Toolkit (GATK) version 3.7-0-gcfedb67 (44) was used to call variants between Illumina reads and the reference. The module ‘HaplotypeCaller with the setting ‘-ploidy 1’ was used to call initial variants between reads and the reference assembly. Then, the module ‘SelectVariants’ was used to select only SNPs. The module ‘VariantFiltration’ was then used to quality filter these SNPs based on the values ‘QD < 2.0’, ‘AF < 1.0’, ‘FS > 60.0’, ‘MQ < 40.0’, ‘MQRankSum < −12.5’ and ‘ReadPosRankSum < −8.0’; the filter flag ‘snp_filter’ was applied to the resulting variant call format (VCF) file.

Following this, the latter two procedures were followed for InDel polymorphisms derived from the initial HaplotypeCaller step. For the ‘SelectVariants’ step considering InDels, the filtering parameters ‘QD < 2.0’, ‘AD < 1.0’, ‘FS > 200.0’, ‘ReadPosRankSum < −20.0’ were used. The filter tag ‘indel_filter’ was applied to the resulting VCF file. The two VCF files and the original VCF file produced by HaplotypeCaller were then merged using the ‘CombineVariants’ module. The settings for this module were ‘-genotypeMergeOptions PRIORITIZE -priority a,b’.

The modules ‘BaseRecalibrator’ and ‘PrintReads’ were then used in conjunction with the merged VCF to recalibrate the BAM alignment. The merged VCF contained higher confidence SNPs and InDels that had passed the quality filtering steps specified in the previous paragraph. HaplotypeCaller was subsequently used, with the settings ‘-ploidy 1’ and ‘--emitRefConfidence GVCF’ to generate a genomic VCF (GVCF) file using the recalibrated BAM.

The GVCF files produced using the recalibrated BAM file were merged using the GATK module ‘GentypeGVCFs’. Two merged VCFs were created based on two groups of individuals, isolates from from Europe and North America, and six isolates from Australia and Morocco. A further merged VCF containing all isolates was also created for analysis of population structure. The joint genotyped VCFs were then hard-filtered again using the GATK module ‘VariantFiltration’. Genotypes with a depth of less than 10 or more than 150 and a GQ score of < 40 were removed. SNPs and InDels that passed this filtering step were retained with the GATK module ‘SelectVariants’. The maximum depth filter of 150 was used to remove repeat-induced alignments and was chosen because it was at the upper-end of the bell curve for coverage in all isolates and below the long tail of repeat-induced coverage (Supplementary figure 2).

### Construction of a whole genome phylogenetic network

The *de novo* assembled genomes produced using the A5 pipeline were used to construct a SNP based distance matrix using Andi version 0.9.6.1 (45). From the distance matrix produced by Andi, Neighbor-net network (46) was constructed using splitstree version 4 (47). A K-mer based distance matrix was also generated using Mash version 2.0 (48). This distance matrix was also used to build a Neighbor-net network using splitstree.

### Principal component and population structure analysis of genotypic variation and analysis of linkage disequilibrium decay

The filtered VCF file containing information from all isolates produced in the previous section ‘Variant calling and quality filtering’ was subjected to a principal component analysis (PCA) using the R Bioconductor package ‘SNPRelate’ version 1.4.2. First, SNPs in strong linkage disequilibrium were filtered using PLINK version 1.90b4.1 (49). To do this, the VCF file was converted into a PED file using VCFTools version 0.1.15 (50). This file was then filtered with PLINK using the settings ‘--indep-pairwise 50 5 0.5’. The coordinates of the resulting SNPs were extracted from the PED file using Awk and then passed to VCFTools to use as a filter with the ‘--positions’ argument. Then, in R, the function ‘snpgdsPCA’ was used with the setting ‘autosome.only=FALSE’ to perform the PCA. Eigenvectors one and two were plotted on the x and y axes, respectively.

Population structure was further assessed using ADMIXTURE version 1.3.0 (51). For this analysis, the PLINK file that had been filtered for SNPs showing strong linkage disequilibrium was used. ADMIXTURE was run for K=1‥10 with the option ‘--cv=10’ for cross-validation.

To calculate linkage disequilibrium decay, a random 30 % of the SNPs for each group were selected using the GATK module ‘SelectVariants’ with the settings ‘-fraction 0.3’, ‘--excludeFiltered’, and ‘--nonDeterministicRandomSeed’. Pairwise linkage disequilibrium was calculated between all SNPs between 1000 and 999999 bp apart using PLINK with the settings ‘--allow-extra-chr’, ‘--r2’, ‘--ld-window-r2 0’, ‘--ld-window-kb 1000’, ‘--ld-window 999999’. The mean value of linkage disequilibrium across 10 Kb sliding windows from any given SNP was then calculated. Loess curves were fit to the mean linkage disequilibrium decay values for the sliding windows with the R function ‘loess’. To determine the approximate distance required to reach 50 % linkage disequilibrium decay, the window start of the closest point above 50 % of the maximum linkage disequilibrium was used.

### Determination of polymorphism effects on coding sequences

The software package SNPEff version 4.3i (52) was used with default settings to determine what coding sequences were affected by SNPs in the filtered VCF from the previous section ‘Variant calling and quality filtering’. The GFF3 file containing reference coding sequences considered in this analysis was downloaded from NCBI (Bioproject number: PRJNA348385). The SNP classifications obtained from this analysis were ‘LOW’, ‘MODERATE’ and ‘HIGH’. These correspond to synonymous mutations with a theoretically lower effect on coding sequences, non-synonymous mutations that may change amino acid sequence without causing major disruption of protein function and highly disruptive non-synonymous mutations that cause alterations such as frame shifts or premature stop codons.

To identify possible sites of transposon insertions into genes, RetroSeq version 1.41 (53) was used with default settings. Read mappings in BAM format from the previous section ‘Mapping of Illumina reads to the reference genome’ were supplied to RetroSeq with the Repet repeat annotation previously generated by Derbyshire et al. (23). To identify genes that were not present in an isolate, the same BAM files and GFF3 file were used to identify genes that did not exhibit any read coverage. Together, genes containing transposon insertions or lacking read coverage were added to the list of genes with ‘high’ impact polymorphisms.

### Re-analysis of published RNASeq data

To determine whether genes associated with selective sweeps exhibited expression profiles consistent with a role in plant infection, an RNA sequencing time course was analysed. The RNA sequencing (RNASeq) data used in this study have been published previously (23,54). In Derbyshire et al. (23), these data were analysed in relation to the version two gene sequences for *S. sclerotiorum*, though only expression data for predicted effectors were presented. In the current study the R Bioconductor package ‘edgeR’ version 3.12.1 (55) was used to re-analyse the RNASeq data and determine significant changes in transcript abundance *in planta* relative to during growth on PDA. The samples included in the dataset (as per the previous publications) were PDA, one, three, six, 12, 24 and 48 hours post inoculation (HPI). Samples were replicated three times and the isolate was the reference isolate, 1980.

Reads were first mapped to the reference genome using TopHat version 2.1.0 (56). The reference gene regions were converted from GFF3 to GTF format using the TopHat utility script ‘gffread’ and specified to TopHat. Then, the function ‘featureCounts’ from the R Bioconductor package ‘Rsubread’ version 1.20.6 was used to generate a feature counts table for each of the gene sequences in the reference genome set based on the GTF feature file and the BAM file. The settings for this function were ‘isGTFAnnotation=TRUE’, ‘requireBothEndsMapped=TRUE’, ‘GTF.featureType=”mRNA”’, ‘GTF.attrType=”gene_name”’ and ‘isPairedEnd=TRUE’. The function ‘DGEList’ from edgeR was used to create a differential gene expression list. Counts per million were derived from raw counts using the function ‘cpm’ from edgeR. Genes were excluded from the dataset if they exhibited less than a total of three counts per million across all replicates.

Next, the functions ‘calcNormFactors’ and ‘estimateDisp’ from edgeR were used to calculate normalisation factors and estimate dispersion, respectively. A generalised linear model (GLM) was then fit using the function ‘glmFit’ from edgeR. The experimental design considered the *in vitro* samples as the reference set and each timepoint was considered relative to during growth *in vitro*. Likelihood ratio tests were used to test each time point against the reference set using the function ‘glmLRT’ from edgeR. The P values from these tests were then adjusted using the Benjamini Hochberg correction (57) for multiple testing with the R core function ‘p.adjust’ and the setting ‘method=”BH”’. Genes were considered significantly differentially expressed *in planta* relative to *in vitro* if they had a p adjusted value of below 0.05.

### Modelling of demographic history of *Sclerotinia sclerotiorum* samples

Dadi version 1.7.0 (58) was used to model demographic history of two *S. sclerotiorum* samples representing two genotypic groups referred to as ‘population 1’ and ‘population 2’ henceforth. These groups were decided on based on the ADMIXTURE, PCA and neighbor-net trees described in a previous section. Population 1 consisted of 11 isolates from Europe and North America and population 2 consisted of five isolates from Australia and one isolate from Morocco.

Before conducting the analysis, VCF files were filtered to remove SNPs with effects on coding sequences (identified using SNPEff) using negative regex expression searches with ‘grep’. The filtered files were then polarised using information from a VCF file, produced using the GATK module ‘HaplotypeCaller’ with the setting ‘-emiRefConfidence ALL’, that contains variants called between *S. sclerotiorum* and the isolate SsChi, which fell within the clade containing *S. minor* from the interspecies phylogeny detailed in the section ‘*HSP60* phylogeny to assign isolates to the species *Sclerotinia sclerotiorum’*. The Python script ‘easySFS.py’ (available at the time of writing at https://github.com/isaacovercast/easySFS) was then used to convert each VCF into a site frequency spectrum (SFS) formatted file, as specified by Dadi. Then, the four single population models in the Dadi built-in functions ‘bottlegrowth’, ‘growth’, ‘two_epoch’ and ‘three_epoch’ were each optimised using the ‘optimize_log’ function for the SFS dataset. The optimised parameters were then used as starting parameters and randomised 20 times using the ‘perturb_params’ function. The randomised parameters were then reoptimised using the ‘optimize_log’ function. Results of the 20 optimisations based on the perturbed parameters were assessed to determine convergence of parameter sets based on SFS data. The model with the highest log likelihood was used to specify 10000 simulations in MS (59). Theta was derived in each case from the optimised models constructed using Dadi. With each simulation, the same number of samples as were in each population were produced for downstream analyses.

### Detection of selective sweeps

Both the real data in the population 1 and population 2 VCF files and the simulated SFS data were scanned for selective sweeps using SweeD version 3.3.2 (5). For each chromosome, the analysis was run separately on a grid size corresponding to positions approximately 5 Kb apart. These analyses were run on all SNPs in the filtered VCF files produced in the previous section ‘Variant calling and quality filtering’. For the simulated SFS data, SweeD was run for the same number of chromosomes as there were samples for each population (population 1 = 11, population 2 = 6). Simulated chromosomes were scanned on a grid of 800 points and were 4 Mb long. Thus, the MS data were also analysed at approximately 5 Kb intervals. A total of 10000 MS simulations were analysed and the maximum composite likelihood ratio (CLR) value for all combined simulations muliplied by two was used as a threshold for detection of selective sweeps. To find genes associated with these regions, Awk was used to generate a BED formatted file containing coordinates 1 / alpha either side of each putative selective sweep. The calculation 1 / alpha is used to obtain the extent of a selective sweep in Nielsen et al. (2005), where alpha is the strength of positive selection. The Bedtools module ‘intersect’ was then used to determine which genes were within these regions.

For each population, Tajima’s D was calculated across a 5 Kb sliding window using the R package ‘PopGenome’ version 2.2.4. Fixation index (F_ST_) between populations was also calculated across a 5 Kb sliding window using this package. Both of these statistics were calculated for each chromosome individually. Both Tajima’s D and F_ST_ were also calculated for all selective sweeps individually and for the whole genome.

In addition to the CLR test, an F_ST_ outlier test was also performed to detect regions of abnormally high fixation between the two populations. To do this, 1000 random genomic segments of equal size to the selective sweeps identified using the CLR method were generated. SNPs present in each of these random genomic segments were retrieved from the previously generated VCF file using the GATK module ‘SelectVariants’. For all of these random segments, F_ST_ was again calculated using the R package PopGenome. Where a detected selective sweep exhibited an F_ST_ value in the upper 95^th^ percentile of the F_ST_ values from the randomisations, it was classed as having a significantly high F_ST_.

### Analysis of genes overlapping selective sweeps

Candidate genes underlying selective sweeps were identified using the coordinates of the selective sweeps and information on SNP impacts on coding sequences, transposon insertions and potential gene deletions. Genes that exhibited non-synonymous SNPs with a minor allele frequency of less than 20 % were considered possible candidates. This is because, if an allele were the cause of a selective sweep, it would be present in most individuals in the population. The same logic was applied to major impact polymorphisms including those identified by SNPEff and transposon insertions and gene deletions. In this case, however, only genes in which the non-reference allele was present in more than 80 % of individuals were considered. This is because the reference gene model was assumed to be the ancestral state, having been modified by the potentially major disruption of function.

To test whether particular GO terms were overrepresented in genes with such polymorphisms in selective sweeps, a GO term enrichment analysis was performed using the R package ‘topGO’. The gene uinverse consisted of all genes with one or more GO functional annotations derived from the InterProScan analysis performed in Derbyshire et al. (23). The algorithm used in this test was the classic Fisher algorithm. The test for over-representation was also carried out for genes without major impact polymorphisms.

## Results

### Assignment of isolates to the species *S. sclerotiorum*

To confirm that fungal isolates indeed belonged to the species *S. sclerotiorum*, a phylogenetic tree was built using previously published *HSP60* gene sequences in conjunction with sequences from *de novo* assemblies. Based on inference from this tree, two originally selected isolates ‘Ss45’ and ‘SsChi’ were dropped from further analyses as they appeared to lie outside the *S. sclerotiorum* clade. Ss45 was not in any known *Sclerotinia* spp. clades whereas SsChi appeared to be in a clade with known *S. minor* sequences (Supplementary Figure 1).

### Population substructure is associated with geography

Whole genome neighbor-net networks, and ADMIXTURE and principal component analyses were used to test for population substructure among the 25 isolates and the reference strain. The ADMIXTURE analysis showed that the value of K (number of hypothetical ancestral populations) with the lowest cross validation error was two (cross validation error = 1.5571). This grouped isolates into two clusters, consisting of those from North America and Europe and those from Australia and Africa. However, cross validation error did not increase much with K and K values of three, four, five and six (cross validation error = 1.2124, 1.31398, 1.35496 and 1.32306) suggested further population structure within the Australian and African population. For K values of three, four and five there was a consistent grouping of Ss44 from Morocco and five Australian isolates into a single population cluster. For all values of K tested, there was little overlap in ancestral population membership between isolates from North America and Europe and isolates from Africa and Australia (Figure 1a).

**Figure 1.**
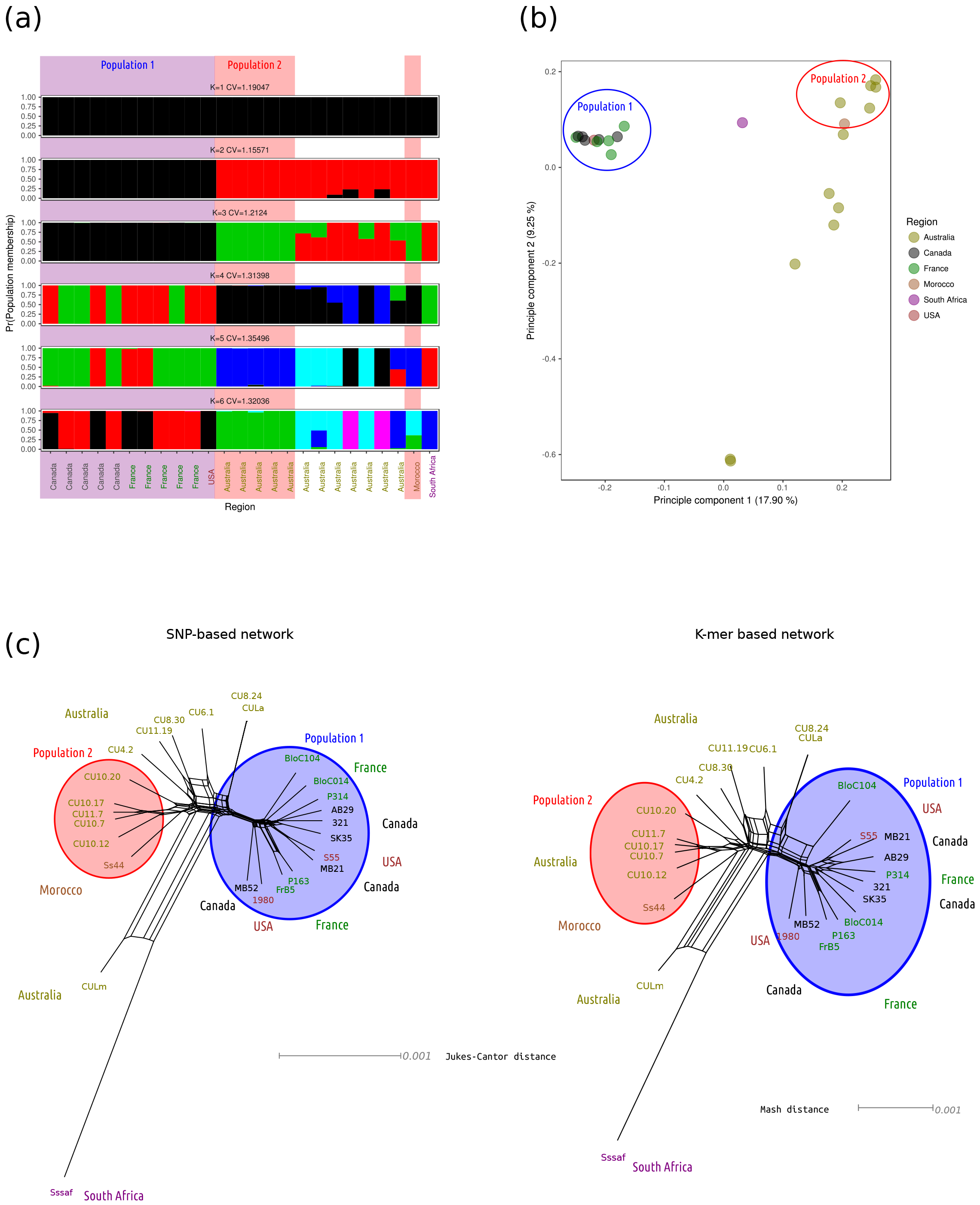
Population structure among the global isolate collection. (a) ADMIXTURE analysis of population structure. Isolates are on the x axis and probability of membership of each hypothetical ancestral population is on the y axis. Bars are coloured according to ancestral populations. Results for K = 1 to K = 6 are presented from top to bottom. Above each plot is the number of K and the cross validation error for that run of the ADMIXTURE algorithm. The two populations used in the rest of the study, population 1 and population 2, are marked with blue and red boxes surrounding plots. (b) Principal component analysis of population structure. The first two principal components are plotted on the x and y axes, respectively. The points are coloured according to region from which isolates were sampled. The two populations, population 1 and population 2, are circled in blue and red circles, respectively. (c) Left: Neighbor-net phylogenetic network built using splitstree with a SNP-based genetic distance matrix generated with Andi. Right: The same as for the left panel but using the mash distance matrix produced by Mash. In both plots, isolates are coloured (and labelled) according to the country from which they were collected. The two populations, population 1 and population 2, are circled with blue and red circles, respectively.

The ADMIXTURE analysis was supported by a PCA, which showed that 17.9 % of variance in genotypic data was explained by differences between isolates from Europe and North America and isolates from Australia and Africa. The second principal component explained 9.25 % of the variance and was associated with differences between population clusters from Australia and Africa. Isolates in the ADMIXTURE population cluster consisting of Ss44 from Morocco and the five Australian isolates grouped together in the PCA (Figure 1b).

The two neighbor-net phylogenetic networks constructed from SNPs and K-mers showed a similar clustering of isolates, but suggested that Sssaf from South Africa was an outlier. All European and North American isolates formed a single cluster in the network and the Australian isolates and Ss44 from Morocco grouped together among the rest of the isolates (Figure 1c). Based on these results, we defined two populations for the selective sweep analyses. These were population 1, which consisted of isolates from North America and Europe, and population 2, which contained Ss44 from Morocco and five of the Australian isolates (Figure 1).

To test whether the two populations exhibited evidence of recombination, linkage disequilibrium (R^2^) was calculated in 10 Kb sliding windows from each SNP in a randomised set of 30 % of the total SNPs. This showed that both populations exhibited evidence of linkage disequilibrium decay with distance (Supplementary Figure 3). Linkage disequilibrium decay reached half its maximum value at approximately 50 Kb in population 1 and 400 Kb in population 2. The maximum R^2^ value in population 1 was 0.51 and the maximum R^2^ value in population 2 was 0.62. This echos the findings of Attanayake et al. (60) who inferred outcrossing in populations of *S. sclerotiorum* based on linkage disequilibrium decay.

### Demographic modelling suggests recent bottlenecks followed by expansions in both populations

The Python module Dadi was used to identify recent demographic events in the two populations of isolates. It was found that a model describing a recent population bottleneck followed by exponential growth fit both of these populations better than any of the other models tested based on log-likelihood (Supplementary Figure 4a-b, Supplementary Table 3). This model took the parameters NuB, NuF, T and probability of misidentification of the ancestral allele (Pr(Ancestral misidentification)). NuB is the ratio of the population size after the bottle neck to the population size before the bottleneck. NuF is the ratio of the current population size to the population size before the bottleneck. T is the time in the past at which the bottleneck occurred. The probability of ancestral misidentification is used to control for errors in ancestral misidentification and create a better estimate for the other parameters. Time is measured in units of 2Ne generations, where Ne is the theoretical ancestral population size.

For population 1, the best fitting parameters for this model were NuB = 0.11, NuF = 0.14, T = 0.46 and Pr(Ancestral misidentification) = 0.09. The log likelihood for this model given the population 1 frequency spectrum was −56.42. For population 2, the best fitting parameters for this model were NuB = 0.075, NuF = 11.049, T = 0.21 and Pr(Ancestral misidentification) = 0.16. The log likelihood for this model given the population 2 frequency spectrum was −18.10 (Supplementary table 3). For both populations, after randomisation and re-optimisation 20 times, model parameters were convergent at higher log likelihoods (Supplementary Figure 4c-d).

### Non-overlapping sweeps appear in both populations

The CLR method first described by Nielsen et al. (4) was used to detect positive selective pressure in population 1 and population 2. The analysis was also run on 10000 MS simulations to determine a threshold value for detection of positive selective pressure. The highest CLR identified for the population 1 simulation was 5.50, and 5.20 for the population 2 simulation. Only CLR values more than double these thresholds were considered for each population. This analysis identified 14 selective sweeps in population 1 and 16 selective sweeps in population 2. These were numbered in order of appearance on chromosome 1 to 16 and are included in the files ‘pop1_sweeps.bed’ and ‘pop2_sweeps.bed’ in Supplementary File 1. Only a single sweep on chromosome 3 was present in both populations (Figure 2a). To expand on these analyses, Tajima’s D was tested for the whole genome and for each selective sweep. Whole genome Tajima’s D was −0.0014 for population 1 and 0.06 population 2. Regions of the genome where selective sweeps occurred generally exhibited lower Tajima’s D than the genome wide figure for both populations (Figure 2b).

**Figure 2.**
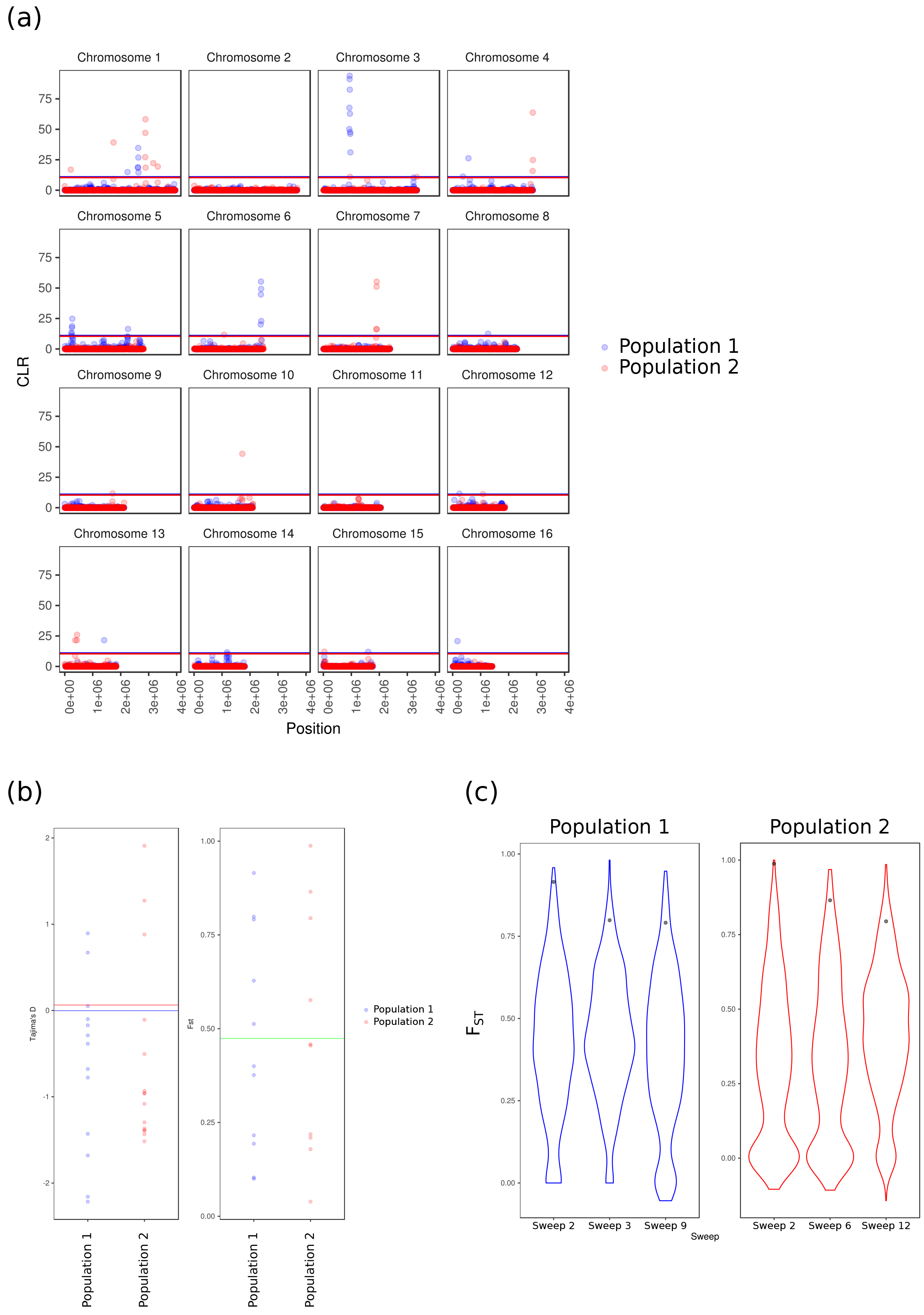
Selective sweeps across the genome of *Sclerotinia sclerotiorum* in isolates from two populations. (a) Each graph represents a different chromosome. On the x axis is chromosome coordinate, and on the y axis is composite likelihood ratio (CLR). The blue points are population 1 and the red points are population 2. The two horizontal lines represent threshold CLR values from data simulated using MS. The red line is the threshold for the population 2, whereas the blue line is the threshold for population 1. (b) Tajima’s D and F_ST_ in selective sweeps. In the graph on the left, Tajima’s D is on the y axis and isolate sample, population 1 or population 2, is on the x axis. The horizontal lines represent genome wide Tajima’s D for population 1 and population 2 isolates. The points represent Tajima’s D for each of the selective sweeps in each of the samples, on the left is population 1 in blue and on the right is population 2 in red. The graph on the right displays F_ST_ on the y axis. Points represent F_ST_ between the two samples for the selective sweeps in population 1 and population 2 and the green horizontal line represents genome wide F_ST_. (c) Selective sweeps with significantly high F_ST_ values. The graph on the left (blue) represents selective sweeps in population 1. The graph on the right (red) represents selective sweeps in population 2. The number of the selective sweep is on the x axis and F_ST_ is on the y axis. Violins represent the distribution of F_ST_ values for the 1000 random genome segments of the same size as the selective sweep. The points represent the F_ST_ values for the selective sweeps.

Since only one of the selective sweeps was present in both populations, F_ST_ between population 1 and population 2 was determined for all selective sweeps and for the whole genome to test the hypothesis of increased fixation of selective sweeps. The genome wide F_ST_ between these two populations was 0.47. Some selective sweeps exhibited a higher F_ST_ than the genome wide figure, but there was a large spread of F_ST_ values among selective sweeps (Figure 2b). Therefore, a randomisation test was used to identify the probability of observing the F_ST_ value of a given selective sweep in a random set of genomic regions of the same size. This showed that there were three selective sweeps in each of the populations that exhibited F_ST_ values that were higher than 95 % of the randomisations (Figure 2c).

### Selective sweeps are enriched for genes with functional terms suggesting roles in regulation of gene expression

To identify genes that might underlie selective sweeps, we performed a GO term enrichment analysis. First, we performed the analysis on all genes that overlapped selective sweeps from either population. This showed that the terms ‘nucleic acid binding’, ‘DNA binding’, ‘intramolecular transferase activity’ and ‘RNA-DNA hybrid ribonuclease activity’ were over-represented among genes within selective sweeps (Figure 3a). Then, we assessed genes with non-synonymous substitutions and major impact polymorphisms such as deletions, disruptive transposon insertions and premature stop codons. In particular, if a gene coincided with a selective sweep, it was deemed to be potentially causal if it exhibited one of these types of polymorphisms in at least 80 % of individuals. Genes that contained polymorphisms of intermediate frequency were not considered as this would suggest appearance of a mutation and partial spread through recombination during or following the selective sweep. Supporting this decision, it was found that among genes in selective sweeps, there was a depletion of alleles of intermediate frequency (Supplementary Figure 5). For non-synonymous polymorphisms, no distinction was made between the ancestral and derived state. However, for major disruptions, the intact reference gene was considered the ancestral state. Both these polymorphic genes and any genes overlapping selective sweeps were subjected to Gene Ontology enrichment analyses.

**Figure 3.**
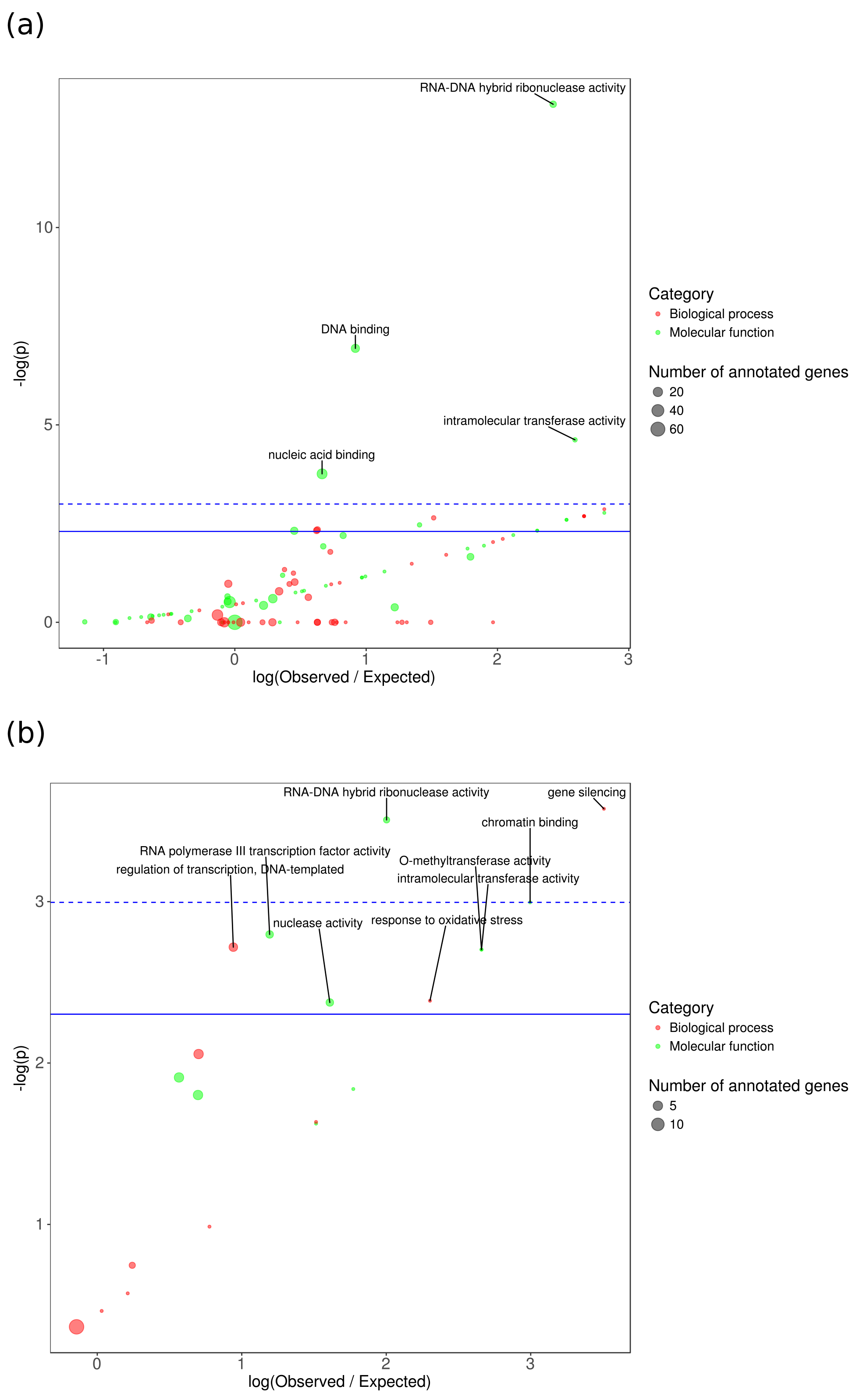
Overrepresentation of GO terms among genes associated with selective sweeps. (a) On the x axis is log(observed / expected) and on the y axis is the −log(p) value of the overrepresentation test (Fisher’s exact test unadjusted). Descriptions of GO terms are indicated with point labels, and the size of the points represents the number of genes containing that GO term that were associated with selective sweeps. (b) Same as for (a) but for genes with high frequency non-synonymous polymorphisms. In both (a) and (b), the solid blue line represents p = 0.1 and the dashed blue line representes p = 0.05.

We found that GO terms for regulatory biological processes such as “chromatin binding”, “O-methyltransferase activity”, “regulation of transcription, DNA-templated”, “gene silencing”, “RNA polymerase III transcription factor activity”, “RNA-DNA hybrid ribonuclease activity” and “response to oxidative stress” had the highest −log(p) values in the test for over-representation among genes with high frequency polymorphisms. Their P values ranged from < 0.05 to 0.099. Among all genes, there were four significantly over-represented GO terms at an alpha of 0.05 (Figure 3b).

Thus, genes with the GO term “RNA-DNA hybrid ribonuclease activity” were overrepresented among all genes overlapping selective sweeps and those that exhibited high frequency polymorphisms. Other GO terms over-represented in all genes overlapping selective sweeps may also be involved in regulatory processes as they suggest binding of DNA.

### The two selective sweeps with the highest CLRs surround polymorphic regulatory and signal transduction genes significantly up-regulated *in planta*

To further investigate selective sweeps in the two samples, the genomic contexts of those with the highest CLRs were considered. The selective sweep with the highest CLR in the population 1 was sweep 3 on chromosome 3, and it partially overlapped with a selective sweep that occurred in the population 2 (Figure 4a). This selective sweep spanned 92,716 bp and exhibited a peak CLR of 93.92. Tajima’s D was decreased in this region to −1.68, as compared with the genome wide figure of −0.0014. F_ST_ in this region was 0.80, which is significantly higher than the genome wide figure of 0.47 (Figure 2c). There were 23 genes that overlapped this region, four of which exhibited non-synonymous polymorphisms that were present in at least 80 % of individuals. Two of these genes were significantly up-regulated from 24 hours post incolation during infection of *B. napus*. These included sscle_03g024870 and sscle_03g025000, which had no predicted functional domains and a methyl transferase domain, respectively (Figure 4c and e).

**Figure 4.**
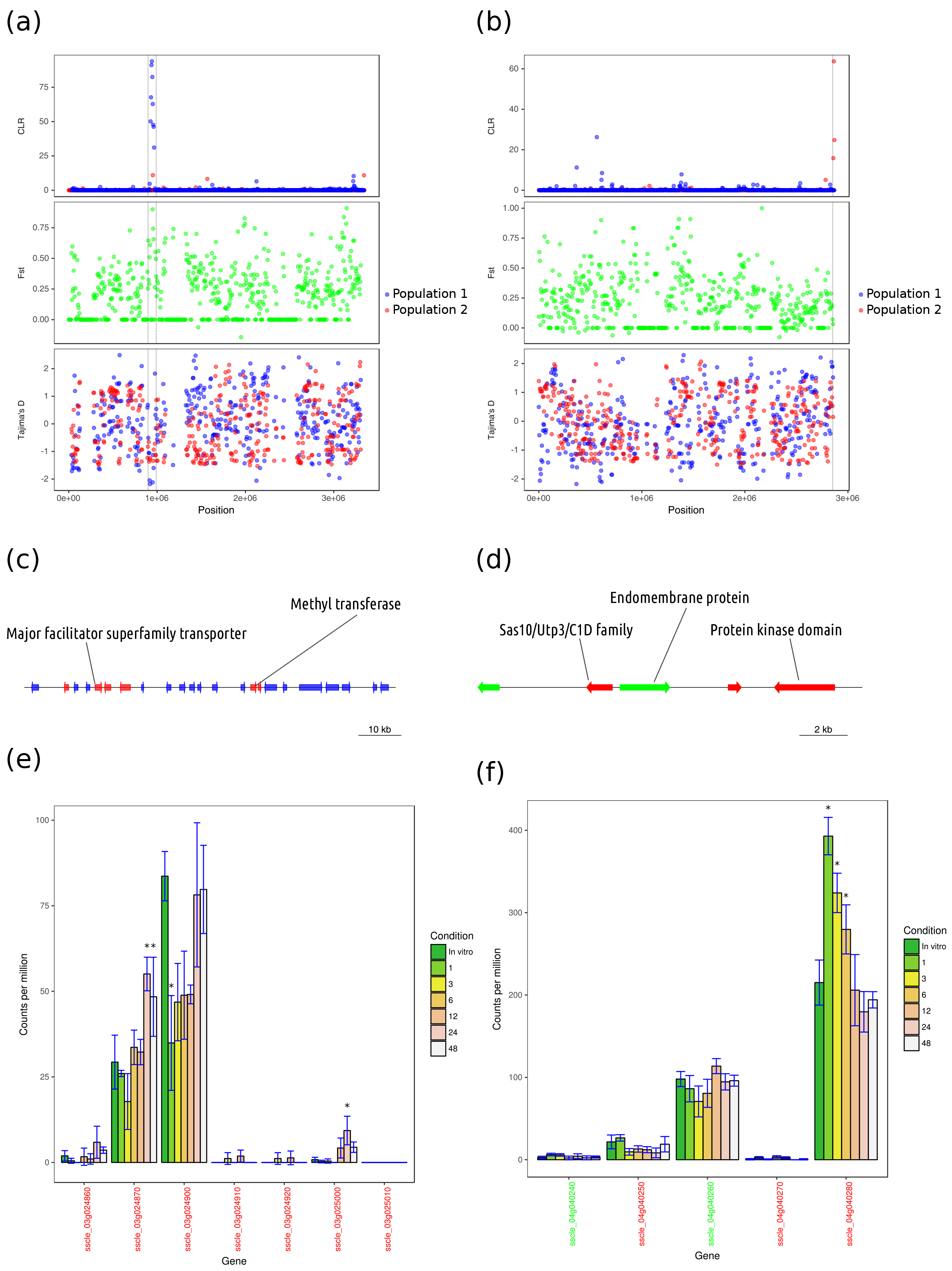
The two selective sweeps with the highest composite likelihood ratios. (a) Sweep 3 on chromosome 3 in population 1. From top to bottom. Top: Composite likelihood ratio (CLR) is on the y axis and genomic position is on the x axis. Blue points represent values for the population 1 and red points represent values for population 2. Middle: F_ST_ is on the y axis and start of each 5 Kb window is on the x axis. Green points represent F_ST_ for all windows. Bottom: Tajima’s D is on the y axis and start of each 5 Kb window is on the x axis. Blue points represent population 1 and red points represent population 2. (b) The same as for (a) but for sweep 8 on chromosome 4 in the population 2. (c) Genes in population 1 sweep 3. Arrows represent genes, with direction representing strand. Blue genes do not exhibit high frequency non-synonymous or high impact polymorphisms. Red genes exhibit high frequency non-synonymous polymorphisms but no high impact polymorphisms. Green genes exhibit high frequency high impact polymorphisms. (d) Same as (c) but for sweep 8 in population 2. (e) Expression of genes in population 1 sweep 3 during infection of the susceptible host *Brassica napus*. Only genes with high frequency non-synonymous or high impact polymorphisms are shown. Counts per million is on the y axis and time in hours post inoculation (HPI) is on the x axis. Error bars represent standard deviation. The y axis tick labels (gene names) are coloured according to whether the gene exhibited a high impact SNP or InDel (red) or was missing completely (green).

The selective sweep with the highest CLR in population 2 was found at the end of chromosome 4. It spanned 21,358 bp and exhibited a peak CLR of 63.70 (Figure 4b). It exhibited a Tajima’s D of −0.11, which was lower than the genome wide figure of 0.064. It exhibited an F_ST_ of 0.18, which was also lower than that of the whole genome. There were only five genes overlapping this selective sweep, and all of them exhibited either high frequency non-synonymous polymorphisms or major disruptions. One of these genes, sscle_04g040280, was significantly up-regulated at one, three and six hours post inoculation of *B. napus*. This gene contained a protein kinase domain (Figure 4d and f).

## Discussion

### Clustering of isolates by geographical origin

Spatial clustering of isolates has been previously demonstrated with genotypic markers such as multilocus haplotypes (MLHs). Carpenter et al. (61) showed that identical MLH fingerprints were shared only between isolates from the same farms in New Zealand. Carbone and Khon (62) suggested divergence between Norwegian populations of *S. sclerotiorum* present in two regions approximately 200 km apart. This was based on a lack of shared fingerprints between the regions, evolution of new fingerprints and increase in size of the intergenic spacer region over a two year period in a specific locale. Sun et al. (63) used random amplification of polymorphic DNA (RAPD) and unweighted pair-group mean analysis (UPGMA) to demonstrate that isolates from Anhui province in China formed a distinct cluster that was distant from another cluster formed from isolates from Poland and Canada. Carbone & Kohn (64) demonstrated divergence between isolates from Wild Buttercup in Norway and populations from different regions of the United States. Haplotypes present in these populations appeared to be exclusive to particular climates, suggesting environmental adaptation. In a further study, Malvárez et al. (65) showed divergence of a Californian population of *S. sclerotiorum* from the previously described North American populations. And, in a more recent study by Clarkson et al. (28), it was found that MLHs were not shared between isolates from the UK, Norway and Australia, possibly suggesting that these isolates belong to distinct populations.

In this study, we show that isolates from various geographically distant North American and European regions form a distinct genotypic cluster (referred to as population 1). The isolates from Australia, South Africa and Morocco formed several distinct clusters that were separate from the North American and European cluster. One of these clusters contained five isolates from Australia and one from Morocco (referred to as population 2). This clustering may suggest adaptation of particular populations of *S. sclerotiorum* to different climates, as was suggested by Carbone et al. (64). Although the various North American and European climates from which isolates in population 1 were sampled are quite different, they all exhibit a break in the season due to a cold or freezing winter and lack of an extensive period of drought (Küppen climate classifiers Cfb and Dfb). This contrasts the climates from which population 2 isolates were sampled, which are characterised by a break in season due to a hot summer with a significant dry period (Köppen climate classifiers Csa and Csb). Fragmentation of the ancestral population into these two populations may have occurred during the last glacial maximum, although this is purely speculation. To identify commonalities between populations from similar climates at the whole genome scale, a further study with more geographically diverse isolates from similar climates is required.

### Linkage disequilibrium decay indicates low levels of outcrossing in population clusters

*S. sclerotiorum* is homothallic, and therefore contains both mating types necessary for successful crossing. It is likely that this favours selfing of the fungus in natural populations, which would lead to extensive clonality. The extent to which outcrossing occurs in nature is unclear, but it has been demonstrated to occur under laboratory conditions. Carbone & Kohn (64) suggested that recombination in contemporary populations of *S. sclerotiorum* is infrequent. This is substantiated by numerous studies in which the same clones have been isolated in successive years, and others in which populations were shown to be composed predominantly of frequently occurring clones. Other studies have suggested that recombination may be more frequent in populations of *S. sclerotiorum*. For example, Attanayake et al. (60) suggested that linkage disequilibrium (LD) decay in seven genotypic groups of *S. sclerotiorum* from diverse geographical regions may be evidence of extensive outcrossing.

In the current study, LD decay in two genotypic clusters suggested some outcrossing between clones. In population 1, which contained five isolates from various parts of Canada, one from the United States and five from France, LD (measured as R^2^) decayed to half its maximum value of 0.52 within 60 Kb. By 150 Kb, LD had decayed to lower than 0. 2. This decay rate is slower than in the selfing plant pathogenic fungus *M. lychnidis dioicae*, in which LD was shown to decay to below 0.2 within 67 Kb (21). This is also slower than the selfing plant *Arabidopsis thaliana*, which exhibits linkage disequilibrium decay to below 0.2 within 50 Kb (66). Population 2, which included five individuals from Australia and one from Morocco, also exhibited LD decay. However, LD decay in this population was even slower than in population 1, reaching half its maximal value at 410 Kb. LD in this population did not decay below 0.3, indicating its alleles were not in complete linkage equilibrium.

Both of these populations thus exhibit evidence of outcrossing and meiotic exchange of genetic material due to decay of LD with genomic distance. However, slow decay in both populations and moderate levels of linkage disequilibrium in population 2 suggests that both sexual and clonal reproduction are prevalent in these populations. The rate of LD decay is expected to increase with time in a sexually reproducing population, dependent on the rate of recombination. The propagation of clonal lineages within a population essentially increases the time between sexual generations for any given individual. Therefore, population 2 could exhibit more clonal reproduction than population 1 or it could be a younger population. As agriculture is a recent enterprise in Australia, populations of *S. sclerotiorum* in this country may indeed have been more recently introduced than populations from North America or Europe.

### Selective sweeps may be evidence of differential selective pressure in two populations

In both populations there was some evidence of outcrossing and meiotic exchange of alleles. Although the samples were likely not drawn from completely panmictic populations, they at least partially met the assumptions of selective sweep scans due to the observed LD decay. To conduct selective sweep scans, we used the method developed by Nielsen et al. (4), which is reasonably robust against assumptions of both underlying demography and recombination rate. We also fit demographic models to the allele frequency data to further control for false signatures of selection. However, discrepancies in parameter estimates for the best fitting models after randomisation and re-optimisation would indicate lack of complete convergence. Despite doubling the highest CLR values from the 10000 simulations, we cannot be completely confident in the appropriate threshold CLR for false discovery of selective sweeps in these populations of *S. sclerotiorum*. The selective sweeps detected in this study should thus serve as a preliminary dataset and may be expanded upon more thoroughly with a larger sample of carefully chosen non-clonal isolates from the same populations.

Interestingly, only one selective sweep appeared in both populations and the rest were in distinct genomic regions. This may suggest that different adaptive alleles have spread in these populations, either ancestrally or due to recent selective pressures. If recombination rate is low within populations, mutation rate of advantageous alleles would also need to be low detect signatures of hard selective sweeps as were identified in the current study. Alternatively, sampling bias due to the relatively small sample sizes may have caused detection of hard sweeps at softly swept loci. Where an allele exhibits multiple unusually frequent haplotypes in a population due to multiple mutations, we may have coincidentally sampled individuals with a single one of these haplotypes.

The different sweeps appeared to be commonly associated with genes involved in processes such as transcriptional or epigenetic regulation, signal transduction and endonucleolytic cleavage. There has been an ongoing debate over the importance of transcription factor evolution in gene regulatory networks. In the past, many favoured the hypothesis that *cis*-regulatory modules are the primary drivers of evolutionary changes in gene expression (67). This is because *cis*-regulatory modules affect the expression of specific genes whereas transcription factors, which bind these modules, often have numerous targets. Changes in transcription factor sequences are thus more likely to be pleiotropic and potentially deleterious (68). However, numerous studies have identified ways in which transcription factors may be able to compartmentalise functions and thus undergo non-synonymous changes without necessarily causing a pleiotropic phenotype. Besides *cis*-regulatory modules, transcription factors often bind multiple different proteins in a modular fashion. Changes in binding affinity for a single protein could lead to more subtle changes in transcription, which might be selected for if they confer fitness advantages. Transcription factors may also undergo alternative splicing, which would again allow for compartmentalisation of function. Particular isoforms may be expressed at particular times and bind particular proteins or *cis*-regulatory modules (68).

Several studies of selective sweeps have given further support for transcription factors as key players in adaptation. In humans, transcription factors involved in processes such as speech and oligodendrocyte myelination have shown evidence of recent positive selection (69,70). In the malaria parasite *Plasmodium vivax* a strong selective sweep encoding a transcription factor was identified. It was suggested that this transcription factor was involved in development of tolerance to antimalarials (71). A further three transcription factors have been identified in *D. melanogaster*, that are associated with strong selective sweeps (72).

Further supporting the hypothesis that gene regulation is an important driver of adaptation, a polymorphic methyltransferase domain-containing gene was associated with the strongest selective sweep in the population 1. This gene was significantly up-regulated during infection of *Brassica napus* at 24 hours post inoculation and it also contained DNA binding domains (data not shown). Methylation of DNA and histone proteins are important regulators of gene expression (73). Several studies have also identified DNA and histone methyltransferases associated with selective sweeps in diverse organisms such as chicken, butterfly and the malaria parasite (71,74,75). Although these studies do not highlight the importance of the methyltransferases they describe - they are, in all three cases, in the supplementary material - it is possible that such genes are subjected positive seletive pressure throughout diverse species. However, without a uniform method of identifying selective sweeps and the genomic loci they affect applied across all species, it is only possible to speculate.

## Conclusion

In this study, we show genotypic clustering of isolates of *S. sclerotiorum* from diverse geographical regions. Climate of origin of isolates hinted at the possibility of population range being dictated by environment. Within two populations, which consisted of representatives from North America and Europe, and from Australia and Morocco, there was evidence of distinct selective sweeps. These selective sweeps may be associated with transcriptional regulation, an observation that has some parallels in previous scans for selective sweeps in other species.

## Acknowledgements

MCD, MD-G and LGK are funded through a bilateral agreement between the Grains Research and Development Corporation of Australia and the Centre for Crop and Disease Management in Curtin University on grant number CUR00023. Part of this work was carried out using the facilities of the Pawsey Supercomputing Centre in Kensington, WA, Australia. MCD would like to personally thank Ellen Elizabeth Derbyshire of Edinburgh University Laboratory for Foundations of Computer Science for guidance and support in data analysis using the diffusion equation employed in Dadi. SR is supported by a starting grant of the European Research Council (ERC StG 336808) and the French Laboratory of Excellence project TULIP (ANR 10 LABX 41; ANR 11 IDEX 0002 02). This project was initiated by Professor Richard Oliver of the Centre for Crop and Disease Management.

## Supplementary figures

**Supplementary Figure 1. Phylogenetic tree assigning isolates to the species *Sclerotinia sclerotiorum*.** The tree is based on heat shock protein 60 *(HSP60)* sequences from other fungi in the *Sclerotinia* genus with the outgroups *Botrytis cinerea* and *Monilinia fructigena*. Node labels are percent support from 100 bootstraps. Tip labels are either isolate names, for the isolates used in this study, or GenBank accessions, for the previously published *HSP60* sequences. To the right of the tree, clades comprising the different *Sclerotinia* species are delineated.

**Supplementary Figure 2. Average genome coverage of Illumina reads for each isolate.** Coverage per variant site is on the x axis and density is on the y axis. The dashed vertical blue lines represent a coverage of 150 x, above which variants were filtered out to remove repeat-induced alignments.

**Supplementary Figure 3. Linkage disequilibrium decay in the two samples of isolates.** Sliding window end position is plotted on the x axis and linkage is plotted on the y axis. Points represent mean linkage for each sliding window for each of the samples with population 1 in blue and population 2 in red. The green dashed lines represent loess curves fitted to the points. The horizontal and vertical red and blue dashed lines represent points where linkage disequilibrium decayed to approximately half its maximal value for each population.

**Supplementary Figure 4. Demographic models describing the observed allele frequency spectra in the two samples.** (a) Model fit for population 1. Top: derived allele frequency is plotted on the x axis and poisson residual is plotted on the y axis. Each point represents deviation of the frequency spectrum produced by the model from the observed data. Bottom: derived allele frequency is plotted on the x axis and the number of sites scaled to the model site frequency spectrum is plotted on the y axis. The data site frequency spectrum is plotted in red, whereas the model frequency spectrum is plotted in blue. (b) The same as for (a) but for population 2. (c) Paramter values after optimisation from random starting parameters for the model that best fit population 1. All tested models are plotted after 20 optimisations. The log likelihood of model fit after optimisation is plotted on the x axis and parameter values are plotted on the y axis. Parameters from all the models are plotted in different colours. The more consistent the parameters are at higher log likelihoods, the better the model fits the data. (d) The same as (c) but for the population 2.

**Supplementary Figure 5. Frequency of non-synonymous and high impact polymorphisms.** (a) On the x axis is the allele frequency, on the y axis is the number of times that polymorphisms at x allele frequency occurred. Data are plotted for all types of considered polymorphisms, including low impact (synonymous), moderate impact (non-synonymous), high impact (disruptive SNPs and InDels), deletion (based on read coverage) and transposon insertion. The blue bars represent population 1 and the red bars represent population 2. (b) The same as for (a) but for polymorphisms within selective sweeps. Many of the alleles at intermediate frequency are reduced from the whole genome distribution, which fits with the drop in allelic diversity that is used to detect selective sweeps.

